# Total recall: episodic memory retrieval, choice, and memory confidence in the rat

**DOI:** 10.1101/2020.12.14.420174

**Authors:** Hannah R. Joo, Hexin Liang, Jason E. Chung, Charlotte Geaghan-Breiner, Jiang Lan Fan, Benjamin P. Nachman, Adam Kepecs, Loren M. Frank

## Abstract

Episodic memory enables recollection of past experiences to guide future behavior. Humans know which memories to trust (high confidence) and which to doubt (low confidence). How memory retrieval, memory confidence, and memory-guided decisions are related, however, is not understood. Additionally, whether animals can assess confidence in episodic memories to guide behavior is unknown. We developed a spatial episodic memory task in which rats were incentivized to gamble their time: betting more following a correct choice yielded greater reward. Rat behavior reflected memory confidence, with higher temporal bets following correct choices. We applied modern machine learning to identify a memory decision variable, and built a generative model of memories evolving over time that accurately predicted both choices and confidence reports. Our results reveal in rats an ability thought to exist exclusively in primates, and introduce a unified model of memory dynamics, retrieval, choice, and confidence.

## Introduction

Animals rely on two sources of information to guide behavior: the external world as it exists in the present, and memory, the internal store of past experience. Perception and memory are both imperfect. Metacognitive monitoring of their possible errors can valuably inform future action, for instance by motivating information seeking prior to decisions, or decreased resource investment afterward^1–6^.

Overall, studies of confidence have focused primarily on information perceived from the external world (*e.g.,* motion detection, odor discrimination), reporting confidence-related behaviors across multiple species including dolphins^7^, non-human primates^8–12^, honey bees^13^, and rats^14,15^. A statistical framework that formally defines confidence and its signatures^14,16,17^ has established a correspondence between statistical confidence in perceptions and the subjective sense of human confidence^18^, and enabled the identification of behavioral and neural confidence markers in species including macaques^10,19^, pre-verbal infants^20^, and rats^21,22^.

By comparison, our understanding of confidence in information retrieved from memory is limited. This is particularly true for confidence in *episodic* memories. This internal computation is thought to correspond to the human sense of confidence in recollections and to go awry in psychiatric conditions^23–26^. Episode memory confidence is also thought to have special implications for consciousness: in Leonard Tulving’s original formulation, three different forms of phenomenal subjective experience (*i.e.*, forms of consciousness) emerge from metacognitive monitoring of the procedural, semantic, and episodic memory systems^27^. Only the third level, autonoetic or ‘self-reflective’ consciousness, is proposed to require — and to imply — conscious self-representation, or self-awareness. It has been argued to exist only in humans^28–30^, and no animal model of episodic metamemory, and thus no demonstration of this state of consciousness or starting point for neurobiological study, has previously been established^31^.

While there is evidence that humans^32^ and primates^12,33–35^ compute memory confidence, studies have focused exclusively on visual recognition memory^33–36^, which can be solved by familiarity rather than true episode recollection^37^. Thus, whether these findings generalize to episodic memories is unclear. Moreover, these studies have not employed explicit models of memory relating choice and confidence, and have not been extended to non-primates. Whether non-primate species compute confidence in memories of *any* type, and how this computation influences behavior, remains unknown.

Here, we developed a behavioral task in rats that enabled quantitative assessment of memory accuracy and confidence for personally experienced events in their temporal and spatial contexts (*i.e.*, *where*, *when*, and *what*, defining features of episodic-like memories in animals^38^). On each trial, rats first made a choice based on information retrieved from memory and were incentivized to then place a bet on whether the choice was correct by waiting for a period of self-determined length. Temporal betting provided a graded confidence report on every trial, improving on task designs that assess only a binary confidence^14,34^, do not allow confidence and choice to be collected in the same trials^8,10,36^, or can only assess confidence on a subset of trials^6,21,22^. This task design enabled collection of thousands of trials from each rat, comprising spatial memory decisions spanning a range of difficulties, each associated with a behavioral confidence report. We found that rats consistently bet more time on correct trials, suggestive of a memory confidence computation. To evaluate this possibility, we constructed a computational model that intuitively unifies memory retrieval, choice, and confidence, and found that it accurately predicts choices and temporal bets.

## Results

### Episodic memory choice and confidence task

We designed a spatial decision task for rats based on episodic memory, augmented with a post-decision wager to assess confidence. Each trial of the episodic memory confidence task requires a binary, memory-guided choice, followed by confidence report (Fig. 1a, b; Supplementary Fig. 1). A randomly selected two of six spatially remote choice ports are cued by a light, and a valid choice is made by entering one of the lit choice ports. The correct choice, or target, is the more temporally remote in the ongoing sequence of visits in the session, while the other, more recently visited port, is the distractor. Next, rats have an option to bet on their choice by remaining at the choice port for a self-determined duration, with the total time spent serving as a bet (Fig. 1b). For correct choices only, longer bets will yield more reward. Importantly, the task takes place in fixed, hour-long sessions, with self-paced trials. Longer temporal bets thus have a higher possible reward payout in the case of a correct choice, but also a higher penalty in the case of an incorrect choice, in the form of the opportunity cost of not initiating a next trial. If rats are able to compute confidence in their memories, to maximize reward over the session they should bet more time on choices based on memories they are more confident in.

**FIG 1.**
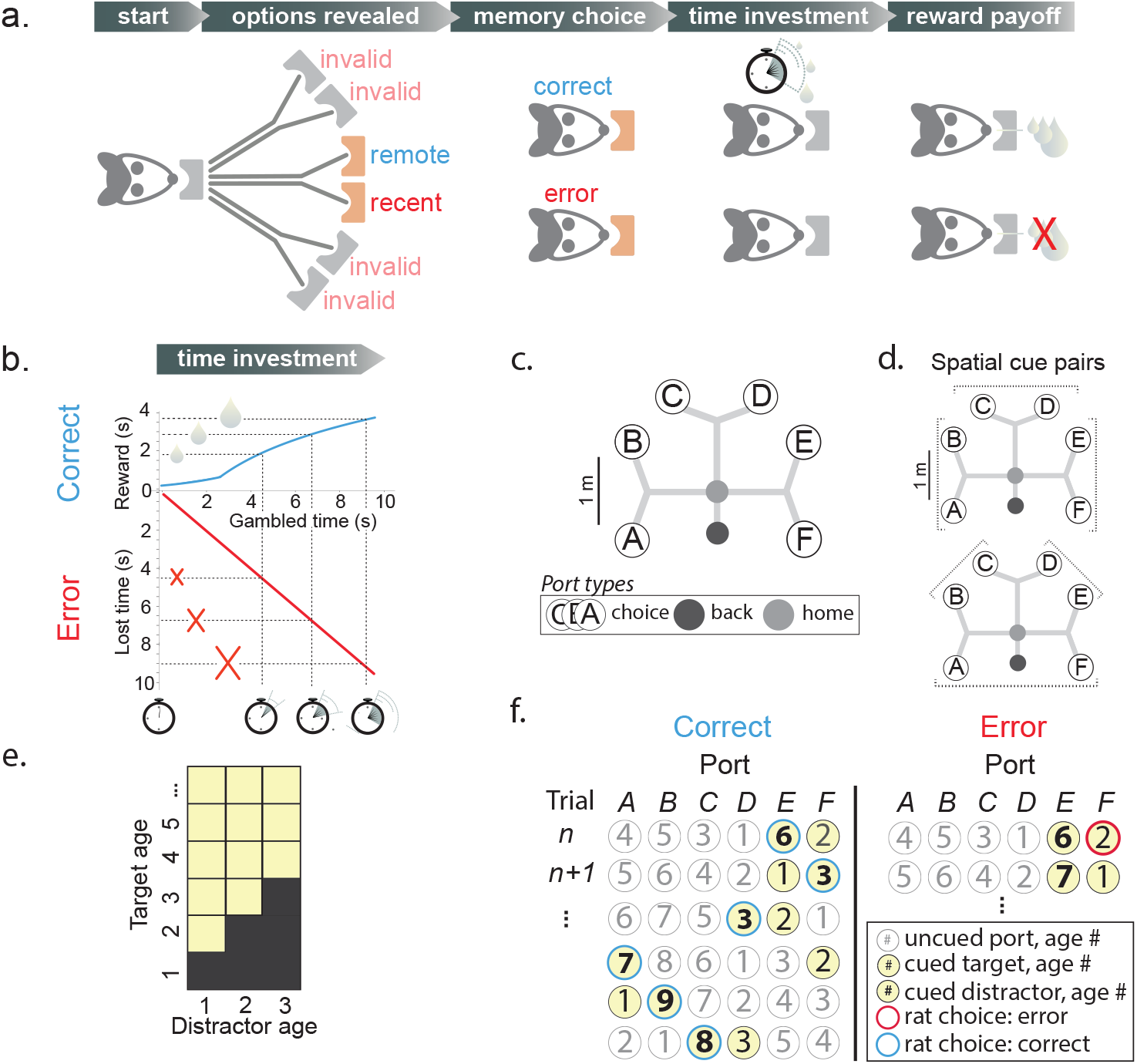
Spatial episodic memory task with time gambling. **(a)** Self-paced trials are initiated by nose-poke at a home port. Two choice port options are cued with a light; four are uncued, invalid options that are not correct. One cued port was visited longer ago in the ongoing visit sequence (*remote*, the *target*) than the other (*recent*, the *distractor*), and is correct. Memory choice is indicated by nose-poke at a port. Time investment: rats gamble on the choice outcome by maintaining the nose-poke position for a self-determined interval. Reward payoff depends, for correct trials only, on gambled time. **(b)** Reward amount (blue) is a function of gambled time, and is received at the choice port. On error trials (red) no reward is received. **(c)** Track geometry showing back (black), home (gray), and choice ports A-F. After leaving choice port, rats receive at back port the same, gamble-dependent reward, completing the trial. **(d)** Cued ports are always adjacent, producing three pairs on the same branch that differ by a stem (top, *stem* trials: AB, CD, EF) and three that differ by both branch and stem (bottom, *branch* trials: BC, DE, FA) trials. **(e)** Distractor ages 1, 2, and 3, with targets older than given distractor, are allowed (yellow). **(f)** Example sequence (top to bottom) of cued ports (yellow) and correct (left, blue outlines) or error (right, red outlines) choices for a range of target (bold number) and distractor (number) ages. After each trial, unvisited port ages increment; last-visited port is set to age 1. Note that trials following error could, but did not usually, present again the same ports.

The reward payoff function was designed to incentivize rats to meaningfully gamble time by countering the possible effects of temporal discounting. Like humans, rats show hyperbolic discounting, preferring smaller rewards sooner to larger rewards later^39,40^, which could counteract the incentive to bet high. Therefore we chose a convex reward payoff function, producing super-linearly increasing reward returns for bets up to 2.2 s (R(t) = 0.27e^0.34(t+0.8)^; Fig. 1b). To discourage excessively long gambled times, we chose a concave payoff function beyond 2.2 s, producing sub-linearly increasing reward returns R(t) = 2.6 × log(0.44 × (t + 0.8)) that delivered 300 μL of reward for the longest typically observed gambled time of 10 s. The briefest registered gamble delivers an approximately 60 μL drop (one minim) of reward, ensuring that rats received an appreciable reward for all correct choices.

The task takes place on a large, branched track, to test memory of episodes occurring at distinct locations as well as times (Fig. 1c). To restrict the number of spatial trial types, the random selection of target and distractor requires that they are always adjacent, resulting in six possible spatial pairs (Fig. 1d). To probe a range of memory difficulties, distractor-target pairs were selected spanning a range of ages (trials since last visit; Fig. 1e). This enabled study of choice accuracy and confidence as a function of how long ago the queried episodes occurred. The distractor age was restricted to 1, 2, or 3. The target age was strictly higher than the distractor age (*e.g.*, for distractor age 1, allowable target ages are 2, 3, 4, *etc.*). Importantly, because distractor-target pairs 1-2, 1-3, and 2-3 are allowable, the task cannot be solved by simply remembering and universally avoiding ports aged 1, 2, and 3. For each rat, the proportion of trials with distractor ages 1, 2, and 3 was approximately one third each, across and within sessions (Supplementary Fig. 2). After each trial, the choice is appended to the ongoing sequence of port visits within the session (Fig. 1f). The correct choice on any given trial therefore depends on the history of actual visits, even if they were errors.

### Rats learn and apply the memory rule with high choice accuracy

Rats took an average of approximately 45 seconds to perform a trial, and thus memory judgments typically related to past experiences on the timescale of minutes. They performed 50 - 100 trials per session and approximately 3000 total trials each, maintaining stable performance accuracy across sessions (see Methods). Choice accuracy was 80.2 ± .04 percent (mean ± s.e.m., *n* = 192 sessions pooled across 4 rats), substantially higher than what could be achieved by a random decision strategy, either across all six choice ports or between the two cued ports (Fig. 2a; Supplementary Fig. 3 a, d, g).

**FIG. 2.**
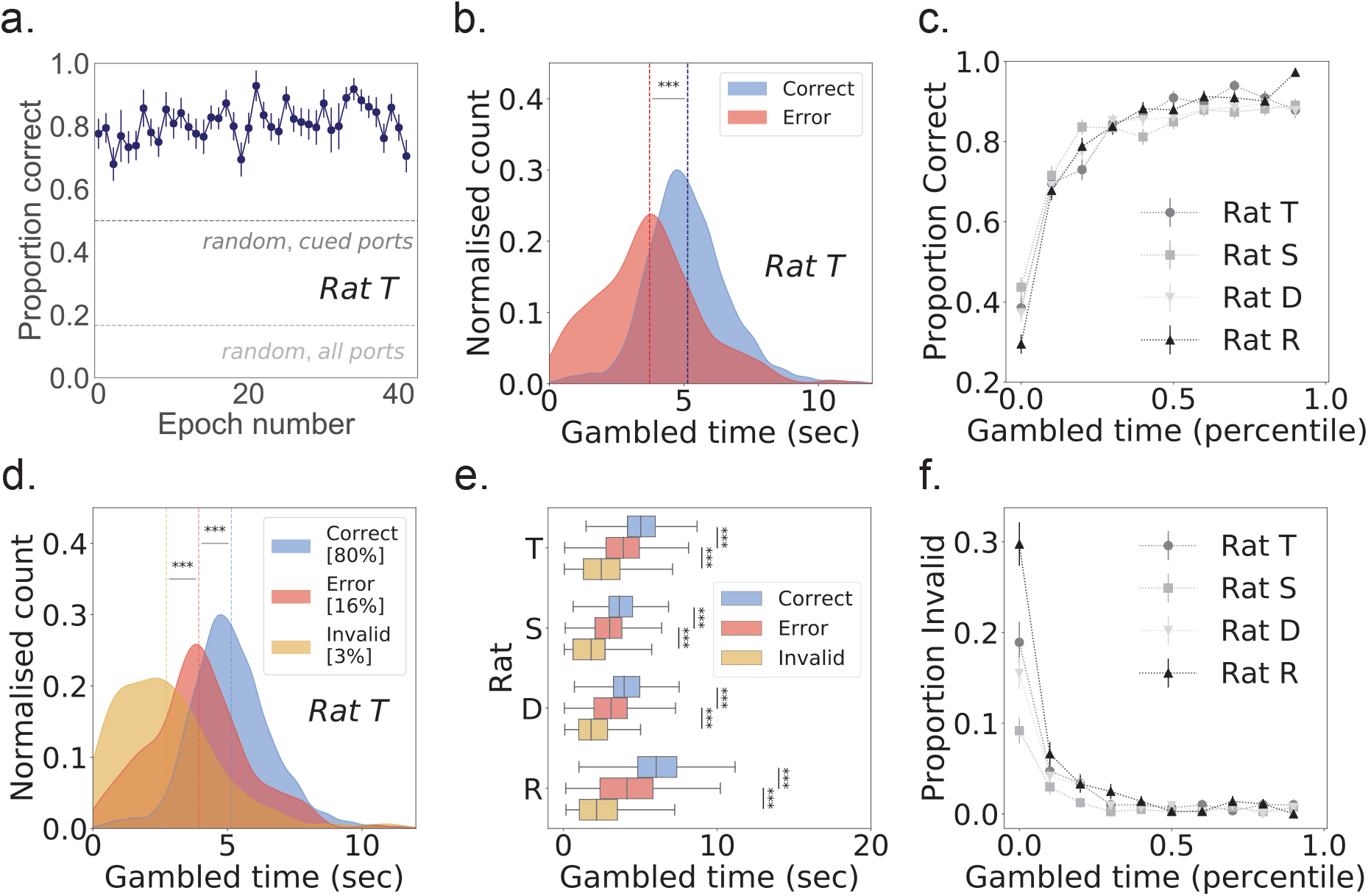
Gambled time predicts choice accuracy. **(a)** Choice accuracy is stable per epoch, as shown for representative rat T at 80.9 ± 0.9%, significantly above random choice between all six ports (light gray line, 17%) or the two cued ports (dark gray line, 50%). **(b)** For representative rat T, average gambled times (dashed vertical lines) were significantly higher for correct (blue) than error choices (red), inclusive over all trials in all epochs (p = 4.8 × 10^−69^). **(c)** For each rat, gambled time (10 percentile bins) predicts choice accuracy, measured as proportion correct. For rats T, S, D, R, *n* trials = 2978, 4111, 4369, 3660. **(d)** For representative rat T, average gambled times (dashed vertical lines) were significantly shorter for invalid choices (yellow) than for errors to the cued port (red; p = 2.5 x 10^−10^). Invalid choices represented the following percentages of total trials: rat T, 3.3%; rat S, 1.7%; rat D, 2.7%; rat R, 4.6%. Excluding invalid choices, average gambled time on correct trials (blue dashed line) is still significantly longer than for errors (red dashed line; p = 6.6 x 10^−48^) **(e)** For all four rats, gambled times for correct trials were significantly higher than error trials (rat S, p = 4.9 x 10^−60^; rat D, p = 5.0 x 10^−81^; rat R, p = 6.5 x 10^−118^.), which were significantly higher than invalid error trials (rat S, p = 2.2 x 10^−9^; rat D, p = 5.6 x 10^−14^; rat R, p = 2.2 x 10^−17^). **(f)** Low gambled times (10 percentile bins) predict a higher proportion of invalid trials for all four rats. All error bars represent s.e.m. and all statistical tests were one-sided rank sum.

Choice accuracy could not be explained by a preference for individual ports or learned port sequences, or by any of a variety of alternative strategies to the true rule (*e.g.*, choose the leftmost of the two cued ports; see Methods; Supplementary Fig. 4). Critically, the stable performance accuracy indicates that temporal bets reflect uncertainty regarding the specific choice rather than uncertainty in the rule itself.

### Temporal bets reflect decision confidence

Rats consistently gambled more time on choices that turned out to be correct (Fig. 2b and Supplementary Fig. 3 b, e, h; average AUC 0.74 ± .03 s.e.m., *n* = 4 rats; for each rat, one-sided rank-sum test *p* ≪ 1 × 10^−5^), and temporal bets predicted overall choice accuracy in a graded manner (Fig. 2c). The difference was striking and consistent across rats: on average, temporal bets were 1.45 ± 0.33 seconds higher for correct than error trials (average ± s.e.m., *n* = 4 rats). Temporal bets were also longer for correct trials considering each port pair separately (Supplementary Fig. 5; for each rat p ≪ 1 × 10^−5^, one-sided rank-sum test). We ruled out the possibility that gambled times were simply a reflection of choice latencies, with longer gambles for decisions that took less time to make. There was only a slight negative correlation (R^2^ < 0.05) despite latency being slightly lower on correct than error trials for each rat (average AUC of 0.55 ± 0.26 s.e.m., *n* = 4 rats; one-sided rank-sum test, *p* < 0.05 for each rat). These results demonstrate that rats can predict choice outcome, consistent with a computation of confidence in their memories.

The rats’ behavior on the occasional visits to one of the two uncued, invalid ports (4.6 ± 0.2 percent of trials, n = 4 rats) also provided evidence for both the knowledge of the rule and a metacognitive assessment of memory choice. The low fraction of these choices indicates that the rats had learned that only cued ports yield reward.

Given that rats understood the task contingencies, their confidence in receiving reward following an invalid choice is predicted to be low, hence little or no time investment in these choices is optimal. Consistent with this prediction, the time gambled on invalid choices was significantly lower than for error trials (Fig. 2d, and Supplementary Fig. 3 c, f, i; average AUC 0.74 ± .01 s.e.m., *n* = 4 rats; each rat, one-sided rank-sum test *p* < 1 × 10^−5^). The fraction of trials that were invalid was highest for the shortest temporal bets, consistent with the possibility that rats understood these trials as exploratory trials with low expected reward (Fig. 2f). Also consistent with this possibility, errors to invalid ports were most common (69.1 ± 3.2 percent, *n* = 4 rats) on distractor age 1 trials (Supplementary Fig. 6), which had the highest proportion correct (Fig. 3a-d), indicating a strategy of selective exploration on easy trials. Hence time bet in invalid trials can be viewed as another form of appropriate metacognitive assessment of memory choice, albeit one that is not formally considered to be decision confidence^41^. Excluding invalid errors, temporal bets were still significantly higher for correct than error trials (Fig. 2d, e, and Supplementary Fig. 3 c, f, i; average AUC 0.71 ± .02 s.e.m., *n* = 4 rats; each rat, one-sided rank-sum test *p* < 1 × 10^−5^).

**FIG. 3.**
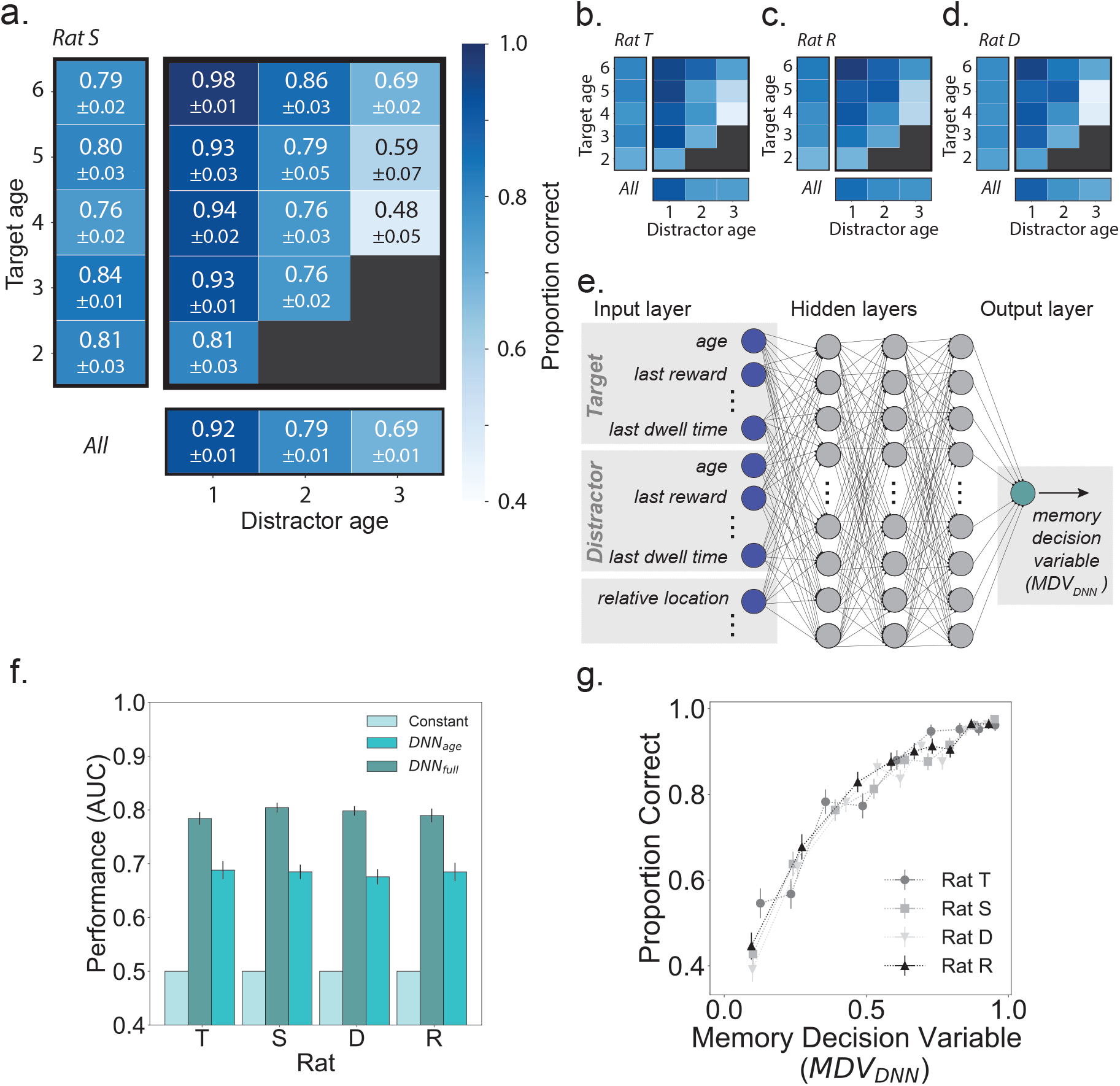
Defining a memory decision variable. **(a-d)** Choice accuracy depends on target and distractor ages. For rats S, T, R, and D, the proportion of correct trials decreases with *distractor age* (columns) and, for a given distractor, increases with target age (rows); marginal performance at left and bottom, respectively. Black boxes indicate trial types not permitted by task logic. **(a)** For Rat S, proportion correct and s.e.m. are annotated. Target ages below 6 are shown, with *n* trials: rat S, 2720; rat T, 2008; rat R, 2499; rat D, 2881. Color bar (a) applies to all four rats. (e) A DNN trained by 5-fold cross-validation for each rat takes as input 20 features, a subset of which are depicted in the input layer (left, dark blue). The DNN has three hidden layers, each with 32 nodes (gray), and outputs a detection statistic related to the probability a trial will be correct, defined as a memory decision variable (MDV_*DNN*_; green). (f) Performance (receiver operating characteristic, area under the curve; ROC AUC) of the DNN trained on the full feature set far exceeded that of a constant model using only the overall proportion correct (*constant*, cyan), as well as that of a model trained on target and distractor ages only (teal). Error bars = s.e.m. (g) For all four rats, a higher MDV_*DNN*_ predicts a higher proportion of correct choices. Horizontal and vertical error bars = s.e.m..

### Choice accuracy depends on memory age and discriminability

What information do rats use to predict choice outcome? We had designed the task to deliver trials spanning a range of difficulties determined by distractor and target ages. If, we hypothesized, choices are based on memory, they should be progressively harder for older targets and distractors^42^. Choices should also be harder for lesser age differences between target and distractor, as episodes that occur closer together in time are more likely to be confused with one another^43^.

Both of these predictions proved to be correct. The average choice accuracies for distractor ages 1, 2, and 3 respectively were 89.5 ± 0.5, 77.7 ± 0.7, and 72.7 ± 0.7 percent (*n* = 192 epochs pooled from 4 rats; Fig. 3a-d). In addition, choice accuracy increased with the age difference between distractor and target when controlling for distractor age (Fig. 3a-d).

### Constructing a synthetic decision variable

Together, these results suggest a memory confidence computation. To evaluate this possibility, we aimed to construct a model of memory confidence, fit to choice accuracy, that would accurately predict confidence and temporal bets as a function of memory discriminability. We therefore had two related goals: first, to characterize the memory discriminability axis for these memory confidence signatures; second, to build a model of memory dynamics as a function of discriminability.

The first step, corresponding to a longstanding challenge in the study of memory confidence, was to identify an appropriate memory discriminability axis, or decision confidence variable^44^. In studies of perceptual confidence the relevant decision variable is typically defined by external task parameters (*e.g.*, motion coherence; odor concentrations) where a simple monotonic relationship between the task parameter and task difficulty can be demonstrated^17^. Alternatively, in the context of value-based decisions, the decision variable is often inferred using a model-based approach that posits a concrete computational model to explain choice behavior^45^.

Neither approach was applicable here: multiple task parameters could potentially influence the rats’ choices, and we are not aware of an existing computational model that could be used to fit the choice behavior.

We therefore sought a model-agnostic approach to derive a synthetic memory decision variable (MDV) that is a scalar summary of the available information that rats could potentially access from memory, with the key property that higher values of the MDV predict higher accuracy. To do so, we trained a deep neural network (DNN) to predict rat choice per-trial based on an exhaustive 20-feature set (Fig. 3e; Methods). We included only those features accessible in memory, not directly observable on the given trial (*e.g.,* previous reward amounts, but not current port identities); hence, a *memory* decision variable. A DNN in particular enabled the agnostic approach we sought: because it is robust to inclusion of redundant and correlated features, an intuitive or model-based feature selection step was not necessary; likewise, selection of interaction terms was not required.

Eighteen of the 20 features were, for each of target and distractor: age in units of trials and time; their last, maximum, and cumulative delivered reward amounts; time since last reward; last and cumulative dwell times; number of trials since any part of its trajectory was last traversed. The final two features were, for the target and distractor, their spatial and temporal (target age – distractor age) trial types. The DNN, trained by five-fold cross-validation for each rat, output a single value, a detection statistic between 0 and 1 that corresponds to a predicted probability that the trial will be correct. As expected, this model outperformed both a model that learned only the overall proportion of correct trials, and a model trained on memory age alone (Fig. 3f). We reasoned that a higher DNN-predicted probability of correct output corresponded to lower trial difficulty, equivalent — because the input features were those available in memory — to *memory discriminability*. Thus, we defined the output of the DNN trained on the full feature set as the MDV*DNN*, with higher values corresponding to memory discriminability and predicting more accurate recall (Fig. 3g). We note that any monotonic function of the inferred MDV will also have the same properties; hence, it is not unique.

### A generative episodic memory model (GEMM)

Having identified a memory discriminability axis, we moved to the second step: building a model of memory dynamics. The MDV provides an index of trial difficulty, but not, in itself, a model of the computation of memory confidence. Moreover, understanding the relationship between the inputs and outputs of a DNN is typically difficult, and may not yield insights into the actual neural computations that underlie animal behavior. We therefore took a more principled approach to develop a computational model of memory confidence, leveraging an understanding of memory phenomena to develop a generative episodic memory model (GEMM) that could predict choice and confidence (gambled time) given an underlying representation of memory.

For this model we focused on memory age, an interpretable and established determinant of memorability that, in our task, independently influenced choice accuracy. We began with a formulation where memory age is represented as a random variable with probability distribution centered on a mental timeline at its time of occurrence. Realizations of this random variable represent specific memory retrievals, corresponding to estimates of how long ago the experience occurred. The distribution’s variance represents mnemonic noise from errors in encoding, consolidation, and/or retrieval. We postulated that (i) these errors accumulate over time such that the memory is less precise, reflected in an increasing variance over time; (ii) the distribution should always take on positive values, as it is not possible to mistakenly retrieve an episode from memory as having occurred in the future, and (iii) an episode should never be completely forgotten.

Given those we developed a mathematical formulation of the model. We define M_α_ as the actual number of trials since the last visit to port α (*i.e.*, the age of that port). Furthermore, we define M_α_’ as the subject’s recollection of the port age. Requirements (ii) and (iii) together specify an asymmetric noise profile with greater spread into preceding than subsequent times. We therefore model M_α_’| M_α_= m_α_ as a lognormal random variable (uppercase symbols denote random variables, while lowercase symbols represent realizations of those random variables). To satisfy requirement (i), the family of lognormal distributions defined by m_α_ = 1, 2, … *n*_*elapsed trials*_ represents the memory’s evolution over time (Fig. 4a). This family of lognormal distributions has a time-dependent mean a_0_m_α_ and a time-dependent standard deviation σ_0_(1 + a_1_m_α_ + a_2_m_α_^2^). We parametrized memory age by elapsed trials and not elapsed time, since a separate regression analysis revealed the number of elapsed trials was a better predictor of choice outcome than elapsed time (Supplementary Fig. 7, see Methods). The separation parameter a_0_ sets the unit increment on the mental timeline that corresponds to one real-life trial; the standard deviation σ_0_ sets the baseline precision of each memory distribution; the coefficients a_1_ and a_2_ set the rate of change for the standard deviation as a second-order polynomial function of its age m_α_, giving it flexibility to increase or decrease as a function of time, though our hypothesis was that it should strictly increase. For a given trial, two ports α and β are cued, with M_α_ < M_β_, corresponding to target and distractor, respectively. Choice (Fig. 4b) is determined by the sign of the difference m_α_’-m_β_’, and confidence by its magnitude, |m_α_’-m_β_’| (Fig. 4c).

**FIG. 4.**
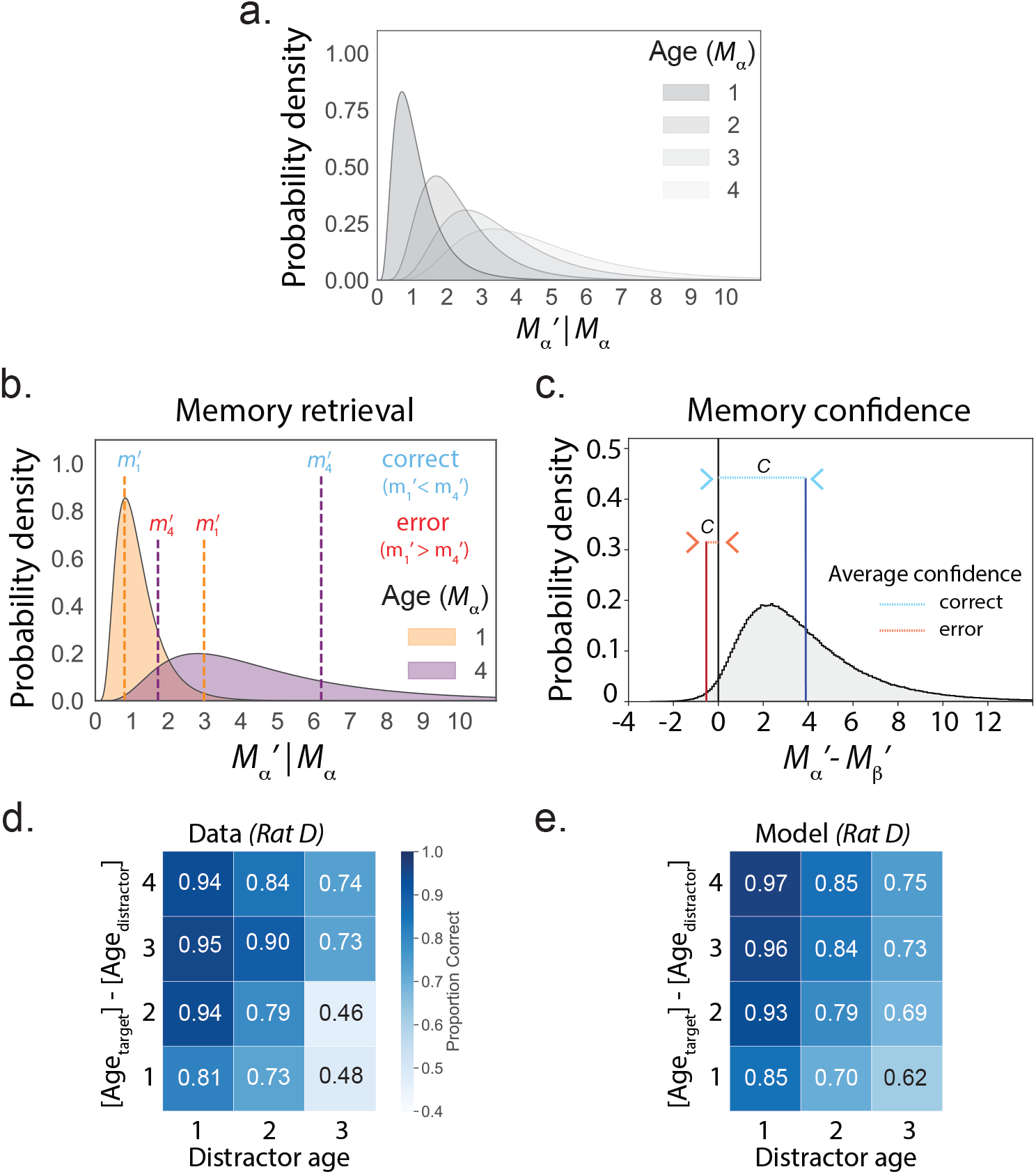
The generative episodic memory model (GEMM). **(a)** Family of lognormal distributions representing the probability density of recalled episode ages M_α_’|M_α_ = m_α_ as the true age m_α_ increments from 1 to 4 for port α. Uppercase symbols denote random variables (*e.g.*, M_α_’,M_α_) while lowercase symbols represent realizations of those random variables (*e.g.*, m_α_’,m_α_). **(b)** Example trial has target port with age m_α_ = 4 and distractor port with age m_β_= 1. A correct (blue) and error (red) realization of the recalled ages for the two ports are shown as vertical dashed lines for the target (purple) and distractor (orange) at values m_α_’ and m_β_’, respectively. **(c)** The probability density of M_α_’ – M_β_’ given M_α_ and M_β_; the area to the right of 0 is the proportion correct for this target-distractor age pair. Confidence (*c*) is computed as | m_α_’ – m_β_’ | and the average confidence is indicated for correct (blue) and error (red) trials. **(d)** Observed choice accuracy across 12 specified trial types, excluding invalid choices. **(e)** Model-predicted choice accuracy across 12 specified trial types, excluding invalid choices. Representative rat D is used for all plots. For rat D, the GEMM uses fitted parameters a_0_ = 1.20, a_1_ = 0.32, a_2_ = 0.38, and σ_0_ = 0.38, for a lognormal distribution with mean a_0_m_α_ and standard deviation σ_0_(1 + a_1_m_α_ + a_2_m_α_^2^). Positive a_1_ and a_2_ define distributions with increasing variance with elapsed trials; σ_0_ ≪ 1 sets a low overlap between neighboring densities, consistent with high observed choice accuracy.

Given that model, we iteratively fit the GEMM parameters for each rat to choice accuracy (Fig. 4d) across trial types based on a χ^2^ metric (see Methods). Based on that fit to memory accuracy (Fig. 4e; Supplementary Fig. 8 a-c, e-g, i-k, m-o), we then generated predictions for memory confidence.

### Embedding the GEMM in data enables prediction of choice and confidence as a function of the MDV

Finally, we combined the MDV and the GEMM to produce a series of confidence tuning curves^22^ to which we could compare behavioral data (Fig. 5). Generating GEMM predictions as a function of the MDV_*DNN*_ enabled the best possible estimates, and ensured our predictions spanned the full range of per-trial memory discriminability. First, for each trial, we input target and distractor age to the previously fitted GEMM to generate a distribution of simulated trial outcomes (correct vs. error) and confidence values (Fig. 5a, *GEMM simulation*). Next, we converted these GEMM-predicted confidence values to gambled times by mapping, for each rat, the inverse cumulative distribution function (CDF) of the observed gambled time distribution (Supplementary Fig. 8 d, h, l, p; see Methods). Note that this mapping has no free parameters. Further, it only considers the full gambled time distribution, not individual trials, and does not separately map correct versus error trials or any other subset of the data, nor does it make assumptions about the match between the mappings of trial outcome to confidence for the model and data. Conceptually, this procedure captures the economic aspect of waiting based on the model, that is, how long the animal is willing to wait given a specific degree of confidence.

**FIG. 5.**
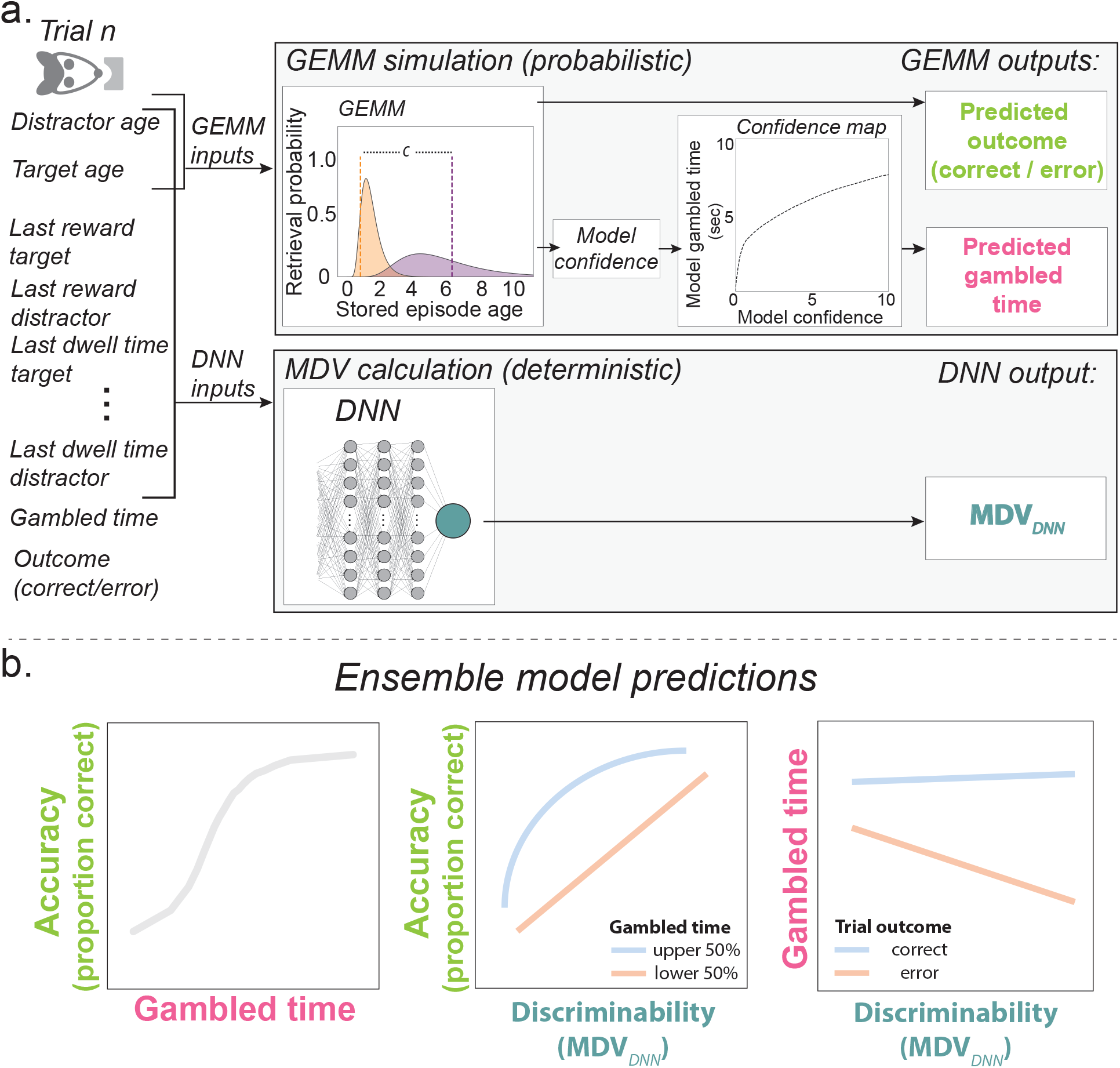
Ensemble model: **A.** For each trial in data, task features (left) include: the 20 features used to calculate the MDV_*DNN*_; gambled time; and trial outcome. A subset of these, the distractor age and target age, are input to the fitted GEMM (top panel) to simulate two GEMM outputs: a predicted trial outcome (correct or error; lime), and a predicted confidence value, which is converted by a monotonic mapping function, shown for representative rat T, to predicted gambled time (pink). The process is repeated *n = 10* times per trial in data to produce a distribution of model-simulated gambled times per observed gambled time, all with the same MDV_*DNN*_ (bottom panel). The MDV_*DNN*_ is calculated from the 20 input features to the trained DNN (green). **B.** The ensemble model makes three signature predictions of memory confidence based on accuracy (lime), gambled time (pink), and the MDV_*DNN*_ (green), as a memory discriminability axis, to which trends in data can be compared (here, representative schematics). Middle panel: blue represents upper half of gambled times; red represents lower half of gambled times. Right panel: blue represents correct trials; red represents error trials.

Every one of these simulated trials has the same MDV_*DNN*_, straightforwardly computed as the DNN output from the 20 input features of the data trial (Fig. 5a, *MDV calculation*). Together, this procedure generated for each trial, (i) a predicted outcome (correct vs. error), (ii) a predicted gambled time, and (iii) a calculated MDV_*DNN*_, which we used to generate three nominal tuning curves for memory confidence based on memory discriminability, temporal bets, and choice accuracy (Fig. 5b). In effect, this procedure generates GEMM-predicted trends for gambled time that are based on all 20 features of the MDV_*DNN*_: although the GEMM only explicitly takes as input distractor and target ages, the GEMM-simulated trials inherit the 20 MDV_*DNN*_ inputs from the data trial they are based on, thereby preserving the covariance structure of the data (*i.e.*, they are embedded in the data, as for hybrid data-simulation models in collider physics)^46^.

### The GEMM accurately predicts memory confidence behavior

We observed a striking match between GEMM predictions and observed behavior. Since all the assumptions of statistical decision confidence also apply to our episodic memory-guided confidence task, we could quantitatively assess the relationship of behavioral confidence reports and GEMM-derived confidence levels by focusing on the established set of comparisons to evaluate confidence as a decision variable^16^. First, a calibration curve makes the intuitive prediction that trials with longer gambled times should have higher choice accuracy (Fig. 6a, d, g, j). Consistent with this prediction, accuracy as a function of gambled time rises for both the model and the data. Second, for any given choice difficulty level (memory discriminability), accuracy should be higher on trials with higher confidence, where more time was gambled. We tested this prediction using a conditioned psychometric curve that divides the data into high and low predicted (GEMM) or actual (data) gambled times. We found consistent and highly similar relationships across data and the model predictions for choice accuracy as a function of memory discriminability (Fig. 6b, e, h, k): longer gambled times predict higher choice accuracy over a range of memorability. Third, for any given trial difficulty level, gambled times should be higher for correct as compared to error trials. Indeed, this “vevaiometric” curve shows consistently higher gambles for correct than error trials over a range of memory discriminabilities in both the model and data (Fig. 6c, f, i, l). For all three signatures and all four rats, the majority of the data points are within two standard deviations of the model, indicating surprisingly accurate fits given the small number of model parameters. This analysis also revealed evidence of an intuitive signature of confidence consistent with the standard model of perceptual decision confidence: the difference in gambled time between correct and error trials is greater for more memorable trials.

**FIG. 6.**
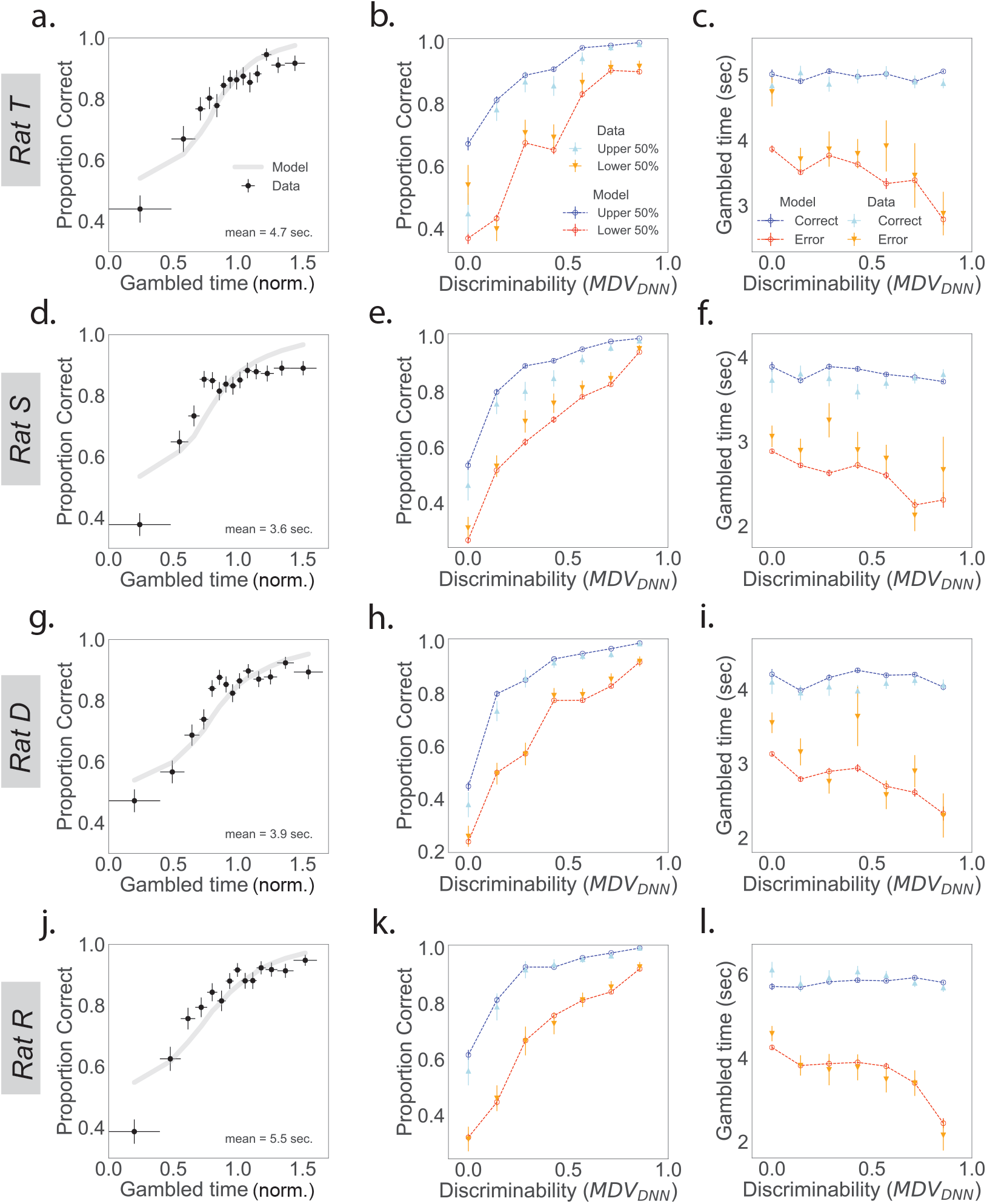
The GEMM predicts trends in memory discriminability, choice, and gambled times in data. GEMM predictions (lines) with data (points) overlaid. **(a, d, g, j)** GEMM-predicted calibration curves (gray lines) for accuracy as a function of mean-normalised gambled time compared to data (black points), for the lowest 14 of *n* = 15 percentile bins. Horizontal bars represent bin widths. **(b, e, h, k)** Conditioned psychometric curve predicted by the GEMM shows proportion correct for upper half (dark blue) versus lower half (red) of gambled times compared to proportion correct in upper half (light blue) versus lower half (orange) in data, each in *n* = 7 percentile bins. **(c, f, i, l)** Vevaiometric curve depicts gambled times predicted by the GEMM for correct (dark blue) and error (red) trials compared to correct (light blue) and error (orange) in data, each in *n* = 7 percentile bins. Vertical error bars represent s.e.m. for all plots.

## Discussion

We designed a spatial episodic memory task to study choice and confidence together. To evaluate memory age as a potential decision and confidence variable, a variety of trials spanned a range of target and distractor ages. We incorporated a novel form of confidence report, time gambling, which was available on every trial. Critically, we found that temporal bets predicted choice accuracy in a graded manner. Our task also allowed us to address the long-standing challenge of defining a memory decision variable (MDV): we trained a deep neural network (DNN) on an exhaustive list of task observables to predict choice accuracy, and interpreted its output detection statistic as defining a synthetic memory difficulty axis or decision variable, the MDV_*DNN*_. Next, we developed a novel generative episodic memory model (GEMM) that posited that the age of memories is represented in the brain as a lognormal distribution that evolves with experiences. We integrated the GEMM and MDV_*DNN*_ in a final model that used the MDV_*DNN*_ to assign a difficulty to each trial and found that across the range of difficulties, GEMM predictions recapitulated the choice and confidence behavior of the animals. These findings provide a demonstration of episodic memory confidence in rats and introduce a simple, interpretable model of the underlying computation.

Studies of learning and memory in animals have typically focused on measures of accuracy, such as time spent freezing in a conditioned context, character of navigation to a hidden platform, or proportion correct^47^. Our results indicate that not only do rats execute behaviors related to representations of past experience, but also that they maintain and can access a representation of confidence related to the retrieval and use of those memories. Rats gambled more time on trials where they had made a correct decision, even though the outcome of the trial was not revealed until after the gambling period ended. Their behavior on trials where they made a choice that was never rewarded (invalid choices) was also consistent with an internal representation of confidence. Rats consistently gambled the shortest times on these trials, consistent with low confidence in a rewarded outcome. In addition, rats rarely made invalid choices and when they did, these tended to be on easy trials (distractor age = 1) with low gambled times. In fact, the lowest gambled times correspond to below-chance accuracy, attributable to a high proportion of invalid trials. This is consistent with an exploration strategy employed selectively on the subset of trials where the true answer is known and “throwing” a trial can therefore ascertain that the optimal strategy is unchanged. Together, these results provide strong support for the hypothesis that rats can access confidence in their memories to guide behavior.

### A model of memory confidence

These findings inspired a two-step approach to understanding the memory confidence computation. First, we inferred a memory decision variable using a DNN with inputs that included variables potentially available to the rats in memory, such as port histories of targets and distractors, and the last reward and last dwell time at each port. The network reached a high degree of prediction accuracy (~80%), outperforming a network based on port age alone.

The output of the network could be interpreted as a decision variable corresponding to trial difficulty. Crucially, in contrast to perceptual ^48^ or value-based ^49^ tasks where the experimenter controls the difficulty of each trial, we did not know how the various elements of each trial would interact to define the difficulty. Thus, this approach has potential to be broadly useful when trial difficulty cannot be established *a priori*.

At the same time, the inclusion of the full 20 parameters, and the complex structure of the network, precluded an immediate understanding of how memory confidence might be computed. We thus focused, in the second, modeling step, on a subset of parameters, specifically target and distractor ages, to design a model to predict choice and memory confidence. Under the GEMM, a few parameters govern the evolution of the underlying lognormal distributions based on known features of episodic memory. Each memory is represented at the time of encoding (t = 0) as a delta function and therefore does not include perceptual noise. At later timepoints, its variance represents mnemonic noise from processes including encoding. The fitted GEMM parameters defined an increasing standard deviation with age, consistent with the understanding that memories become less precise over time and that when a memory is retrieved for consolidation or use it can become labile again^50^. The instantiation of known features of memory, combined with flexibility in their evolution parameters, means that the GEMM could be applied to other tasks as well (*e.g.,* recognition memory tasks). The GEMM could also be adapted to describe memory dynamics not just for the recalled time of episode occurrence, but for the details of those episodes themselves. For instance, remembered places or faces might blur over time into similar ones with an increasing, time-dependent variance as in the GEMM and, for complex stimuli such as these, its asymmetric noise profile as well.

Although the GEMM describes memory dynamics as a function of age only, the other MDV_*DNN*_ features are taken into account by its embedding in the data in the full ensemble model. Importantly, the GEMM was fit only on the trial outcomes, and it nevertheless provided remarkably accurate predictions of gambled time. The key feature that enabled those predictions is the asymmetric noise profile, which is motivated by known dynamics of memory decay. Critically, this differentiates it from the fixed-variance Gaussian noise profile that is typically used in perceptual discrimination and that is implicit in many models of recognition memory ^37,51^. We found that behavioral data for all four rats were well described by the GEMM, which predicted confidence tuning curves similar to those of statistical confidence with one exception: only the GEMM explained the relatively constant gambled times for correct trials a function of memory discriminability seen on the vevaiometric curve.

Importantly, achieving relatively constant gambled times for correct trials is non-trivial. An alternative approach to generating confidence predictions from the GEMM would be to use the MDV_*DNN*_ as a decision axis, assume that on each trial the memory decision and memory confidence are both determined based on a simple noise profile (*e.g.,* Gaussian noise with fixed variance), and from this predict gambled time^21,22^ (Lak et al, 2014; Masset et al, 2020). Such a model would predict that confidence increases for correct trials and decreases for error trials as a function of discriminability^16^. However, in our data, gambled times do not increase with the ease of the decision. This emphasizes the importance of identifying the appropriate noise distribution producing choice variability. Further, even an approach combining the MDV_*DNN*_ with a non-Gaussian, non-fixed noise profile would have limited utility: reliance on the DNN would obscure the minimal set of variables that were sufficient to construct the GEMM. As such, this approach would make it more difficult to make specific predictions about potential representational substrates of a memory confidence calculation in the brain.

### A framework for understanding a memory confidence computation

Our findings constitute evidence of an ability to compute and act on a representation of memory confidence in a non-primate consistent with the notion of autonoetic consciousness, opening the door to a mechanistic understanding of the underlying neural processes. Previously, human and non-human primate studies have identified spiking activity reflective of recognition memory confidence^33,34,36,52^, but the relationship of this activity to the neural activity underlying memory itself is unknown, and difficult to quantitatively model in the absence of clearly defined memory decision variables (*i.e.*, discriminability axes). A possibility suggested by the GEMM is that a memory confidence computation receives input from the circuitry of memory retrieval itself. Indeed, lognormal distributions are commonly observed in the hippocampus, a brain structure critical for episodic memory processes^53^. The GEMM suggests such questions as whether memory age might be encoded as a lognormal distribution of firing rates during memory retrieval^54^.

Information from the hippocampus and related structures might provide input to a confidence computation, but current evidence suggests this computation most likely occurs outside the hippocampus. In rats, many studies have localized perceptual decision confidence, but not the decision itself, to the lateral orbitofrontal cortex (lOFC)^21^, including, most recently, modality-general perceptual confidence representations^22^. It is possible that the same lOFC neurons could also represent memory confidence, particularly since memory itself is multi-modal. We hypothesize that perceptual and memory confidence are integrated in decision-making, which optimally depends on evidence from varied sources, on long timescales. Prior to this step, different forms of memory confidence may be computed differently, by different circuits^35^.

While aberrant confidence computations have been proposed to account for a variety of psychiatric symptoms including hallucinations, these ideas are based on models of perception^26,55,56^. In contrast, the study of memory has not had the requisite behavioral tasks, animal models, or theoretical framework for understanding memory, confidence, and their roles in disease. A deeper understanding of memory confidence has potentially broad applications, for instance in judging the credibility of eyewitness testimony (*e.g.,* in the 2018 Kavanaugh hearings)^57^. Ultimately, a complete description of how memory guides behavior should include confidence. Toward such an understanding of total recall, we propose as possible starting points trial-based memory tasks, DNN-derived synthetic decision variables, and the GEMM as a first phenomenological account.

## Methods

### Behavioral training and task

All procedures followed the guidelines from the University of California San Francisco Institutional Animal Care and Use Committee and US National Institutes of Health. Male Long-Evans hooded rats were trained to perform an episodic memory task with time gambling for liquid reward. Behavioral testing was controlled by custom software written in Python using data acquisition hardware (Trodes ECU, SpikeGadgets LLC) to record rat pokes and unpokes at the ports and to control reward delivery.

#### Habituation

Three cohorts of Long Evans male rats (3-4 months old, 450-600g, 6-8 rats per cohort) were habituated to daily handling for a week and to hand-delivered liquid food reward (evaporated milk plus 5 percent sucrose) from a syringe in the home cage for three days.

#### Stage I: Raised linear track plus delayed reward

Animals were then food deprived to 85-90 percent of their baseline weight and pre-trained on a raised linear track for 3-4 days, 2-3 epochs/day, 10 mins/epoch. A port was located at each end of the track, equipped with an LED light and an IR beam, to detect entry and exit from the port. Each port could automatically deliver reward, which was available for only a specified length of time as it flowed through the port at a rate of 0.17mL/sec to a drainage outlet and did not remain in the port. A variable delay τ between nose-poke and reward delivery was drawn from an exponential distribution, which was gradually incremented from τ = 0.2 − 0.5 seconds to τ = 1 − 8 seconds. Only one port was cued by a light on each trial. After nose-poke detection, the light went out and reward was delivered. The two ports were lit alternatingly over the course of the epoch. Rats learned to run back and forth on the track to visit the currently lit port and to wait for the delayed reward. From each cohort, 2-3 rats with the highest accuracy and speed were selected for training on the episodic memory confidence task.

#### Stage II: Full episodic confidence task sequence with experimenter-delayed reward

In Stage II, rats learned the basic task structure (Supplementary Fig. 1), but with only one cue lit per trial and a pseudo-gambled time determined by the experimenter. The track has eight ports in total: one home port at the center, one back port, six choice ports at each end of six branches. As in Stage I, each port could be cued with a light and deliver liquid milk reward. Each epoch was of a fixed length per animal, during which trials were self-paced. The lit cue corresponded to the target selected by the same code as in the final task logic; lighting of the distractor port was suppressed. The sequence of visits within a trial was: home port light on; rat pokes at home port for a small fixed reward (350 ms); home port light off; after a variable cue delay, one choice port light on; rat pokes lit choice port; choice port light off and port delivers initial reward (350 ms) and, after a variable, experimenter-controlled reward delay, a wait-dependent reward; back port light on; rat pokes back port; back port light off and port delivers back reward. Choice accuracy was measured as the percentage of trials for which the rat visited the lit choice well.

The cue delay was introduced to jitter the events of each trial relative to every other trial, to control for across-trial temporal correlations between behavioral and neural events. To train rats to wait for the cue lights to come on, the cue delay was gradually increased from range [0.2, 0.5] to [0.5, 2.0] seconds. Initially, the back port delivered the same reward amount as the wait-dependent reward regardless of trial outcome, which encouraged the animals to solidify knowledge of the port visit sequence (*i.e.*, to not skip the back port). After three epochs, back port reward was only delivered on correct trials. The reward delay was determined by sampling from an exponential distribution with rate parameter λ = 1/2, accepting only samples that were between 1-3 at the start of this training phase and 2-10 by the end, with a wait-dependent reward amount that increased accordingly, to allow rats to learn that a longer period spent nose-poked in the port would result in a larger reward.

#### Stage III: From delayed reward to gambling

After rats were consistently performing at above 80 percent choice accuracy and waiting for the full reward delay, the initial reward was omitted. Once rats were able to wait for the majority of the reward delays (6-10s), the switch was made to gambling logic. In the gambling logic of the final task, rats voluntarily reported the time they were willing to wait for a potential reward. The gambled time began at the time of nose-poke in the choice port and ended when rats withdrew from the port. Nose-poke withdrawal was detected with a ‘grace period’ (800 ms for rats T, S, D; 700ms for rat R in final behavior, calibrated based on how quickly each rat moved) to allow for small head movements during the gambling period: rats were only declared to have ended the gambling period after a grace period had passed between the port’s IR beam re-forming (un-poke) and being broke again (re-poke). After gambled times were observed to be stable across at least three epochs, the distractor cue was introduced alongside the target cue, starting with distractor age 1. Distractors age 2 and 3 were introduced when choice accuracy was approximately 80 percent and stable.

#### Stage IV: Data collection

Approximately 3000 - 4000 trials were collected from each of four rats. Each rat had a typical length of time for which he would continuously perform the task, after which he would occasionally perform trials but otherwise sleep or lean off the edge of the track and attempt to eat the milk tubes or CAT6 cables. Epochs shorter than 20 minutes (Rat T, n = 5 excluded epochs), 40 minutes (Rat S, n = 0 excluded epochs, and Rat D, n=2 excluded epochs) or 45 minutes (Rat R, n = 2 excluded epochs) were excluded from final analyses. This resulted in the following epoch and trial counts: From rat T, 2978 epochs over 42 trials; from rat S, 4111 trials over 40 epochs; from rat D, 4369 trials over 61 epochs; from rat R, 3660 trials over 49 epochs. Typically rats ran an average of 350-400 meters per day (the human equivalent of approximately five miles) and consumed 50 mL of sweetened evaporated milk.

#### Parameter setting: distractor and target selection

The selection of distractor and target was random with temporal weighting, to guarantee that trials with distractor ages 1, 2, and 3 were evenly distributed throughout the epoch. During an initialization period, the rat was cued to visit each of the six choice ports in a randomly generated order, establishing a history of visits. After every port was visited at least once, the logic used for selection of the two cued ports on each trial was: from the list of possible port pairs with their ages, for example, the top row of Fig. 1f, [AB(4,5), BC(5,3), CD(3,1), DE(1,6), EF(6,2), FA(2,4)], select candidate pairs for which at least one of the ports has an allowable distractor age (1, 2, or 3), which are [BC(5,3), CD(3,1), DE(1,6), EF(6,2), FA(2,4)] here. If there is more than one candidate pair in this list, remove from it the candidate pairs with distractor ages equal to those presented on the last trial, the penultimate trial, and the trial before that, in that order, until candidate pairs with only one distractor age remain. If there is only one candidate pair in this set, select it as the presented pair. If there is more than one candidate pair in this set, randomly select between them with equal probability. For example, if the last three trials were distractor ages 1, 2, 3 (*N.B.*: regardless of which ports these distractor ages corresponded to), then on the upcoming trial, the candidate pair(s) with distractor age 3, [BC(5,3)], would be removed first, then the candidate pair(s) with distractor age 2, [EF(6,2), FA(2,4)]. The candidate pair(s) with distractor age 1, [CD(3,1), DE(1,6)], would be selected; if there were more than one candidate pair with distractor age 1 remaining, the cued pair would be selected randomly from this set. On every trial, there will necessarily be a candidate pair with distractor age 1. There will not, however, be candidate pairs with distractor ages 2 and 3 on every trial; this can occur in the case of revisits, where the port with distractor age 3 is the same as the port with distractor age 1 (or the age 2 port = the age 1 port, or the age 3 port = the age 2 port = the age 1 port). This selection algorithm has the effect of sampling evenly across distractor types, resulting in approximately 1/3 each per epoch and preventing an alternation sequence from developing.

#### Parameter setting: reward function

The reward function was designed to counter the potential effects of temporal discounting on gambled times. The expected effect of such temporal discounting is that rats would reduce their gambled times to receive a smaller reward sooner rather than waiting for a larger one. This effect may be greater on trials where they are highly confident in their memories and choice, as the option of a smaller reward sooner is more certain. This effect could obscure the difference between gambled times on correct and error trials by inducing a left shift of gambled times on correct trials. To counter this possible effect, the reward amount delivered was a piecewise function of gambled time with a relatively low derivative for the first 2.2 seconds and a relatively high derivative after 2.2 seconds (Fig 1b). On correct trials, for investments less than 2.2 seconds, the length of time for which a sweetened evaporated milk reward was delivered at a constant rate of 0.17 mL/sec was given by R = 0.27e^0.34(t+0.8)^; for investments greater than 2.2 seconds, R = 2.6 × log(0.44 × (t + 0.8)). A ten-second wait, for example, will yield a four-second reward. The desired effect was to bias the rat toward longer gambled times on trials for which he would already have waited at least 2.2 seconds, as he could double the reward amount by waiting just one second longer. If rats were able to access memory confidence, these longer waits should be more common for correct trials, and the reward function could help resolve them from error trials. The non-zero intercept ensured that the rat received an appreciable reward amount (350 ms, 60 μL, equal to approximately one drop, or minim) even for very short waits on correct trials, preventing the development of uncertainty in the memory rule itself following correct trials that resulted in zero reward due to short gambled times. To ensure a high enough number of trials per epoch to sample trial types evenly, we discouraged extremely long gambled times greater than 9.5 seconds by choosing a reward function with a derivative that fell by 9.5 seconds to the level it was prior to 2.2 seconds. Rats took an average of 15 seconds to perform a trial excluding gambled time. With a 9.5 second gambled time and the resulting 4-second reward delivered at both choice and back ports, this yields approximately 30-second trials and our aim of 80 trials per 40-minute epoch.

Rats that performed many trials per epoch with a large spread in gambled times were implanted with hardware for recording neural data. Following a week or more of recovery, behavioral data in the final task were acquired from implanted rats.

### Correlation of choice latency and gambled time

For analysis of correlation between gambled times and latency to choice, outliers with gambled times greater than 10 seconds or latency to choice greater than 20 seconds were excluded, leaving over ninety percent of the data per rat. Linear regression was implemented in SciPy.

### Evaluation of alternative strategies

For each rat, the proportion of times that each port was presented as target versus distractor were compared. Per epoch, these values were rarely above or below 50 percent by greater than 3 percent, and the majority of differences were not statistically significant at p = 0.05 by a t-test for independent samples.

We tested whether there existed an alternative strategy that could better explain the rat’s choices than the true rule, which is to select the least recently visited of the two cued ports. For every trial in every epoch, for each rat, we determined whether the alternative rule would have resulted in the same choice as the one the rat made, or the same choice dictated by the true rule. This resulted in two proportions per epoch for each rat.

### Evaluation of logistic regression and neural network models of choice accuracy

We used a DNN model to predict choice outcome (correct or error) as a function of an exhaustive feature set, or a feature set comprised of target age and distractor age alone. The exhaustive feature set included for each of target and distractor: age in trials and time; their last, maximum, and cumulative delivered reward amounts; time since last reward; last and cumulative dwell times; number of trials since any parts of its trajectory was last traversed. The feature set also included, for the target and distractor, their spatial trial type (branch/stem) and temporal (target age – distractor age) relationships. The features were each standardized to have zero mean and unit variance. The DNNs were feedforward, fully connected networks implemented in Keras using the Tensorflow backend and optimized using Adam. Each network has three hidden layers with 32 nodes each and the rectified linear unit activation. The output of the last layer is a sigmoid and the binary cross-entropy is the loss function. Networks were trained with 200 epochs with early stopping using a patience of 5 epochs. A k = 5-fold training procedure is used whereby 1/k^th^ of the data are withheld for testing, 1/k^th^ are withheld for validation and the rest are used for training. Datasets used for training were subsets of the full dataset for each of rats T, S, D, R (*N =* 2857, 4031, 4246, 3452, respectively) due to the requirement that training trials have data for every feature in the exhaustive set. The trials that comprise each fold are uniformly selected at random. A total of 10 networks are trained for this configuration and the network with the best validation loss is used to evaluate on the test set. The test set is then rotated k times until all data are used for testing. The loss is weighted during training so that the weighted number of instances from the two trial outcomes (*i.e.*, correct or error) are the same.

Logistic regression was implemented in Keras, where it is simply a neural network without any hidden layers.

### Fitting the generative episodic memory model (GEMM) parameters

The GEMM was fit on a subset of distractor-target trial types for which there was enough data, excluding invalid errors. The reduced datasets were 1877, 2593, 2722, and 2284 trials for rats T, S, D, R, respectively. Model parameters a_0_, a_1_, a_2_ and σ_0_ were fit for each rat based on its performance across trial types defined by distractor and target - distractor ages (excluding invalid error trials and target - distractor ages > 4). The probability density of the difference between two lognormal distributions (whose negative density is the error rate) does not have a closed-form analytic solution, so we simulated 10^4^ trials for each trial type within the fit. Each simulated trial generated an m_1_’ and m_0_’, from which we computed an outcome (*correct* or *error*). Across many simulated trials, this returned a predicted error rate pattern across trial types for the current set of parameters.

A χ^2^ metric was used to evaluate model performance and find the best fit parameters:

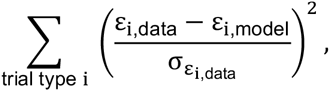

where ε is the error rate and σ_ε_ is the uncertainty in the error rate. The uncertainty σ_ε_ is determined via bootstrapping, accounting for correlations between the number of trials that were incorrect (N_i_) and the total number of trials (N_T_) by modeling each as an independent Poisson random variable and taking the standard deviation of N_i_/(N_i_ + N_c_) over 100,000 simulated trials. We use the Nelder-Mead method with 200 maximum iterations as implemented in SCIPY, minimizing the χ^2^ fit to error rates across trial types. Then, using these parameters, we generated the distributions corresponding to each episode memory and sampled from each 100,000 times to generate target memories, distractor memories, the outcome of the trial (correct/error) and a confidence (absolute value of the difference between target and distractor).

### Mapping GEMM-predicted confidence to gambled time

To convert the simulated confidence values to invested times, we mapped the confidence (C) probability density onto the probability density of the rat’s invested times (T). Let F(x) = Pr(C ≤ x) be the cumulative distribution function (CDF) for C and G(x) = Pr(T < x) be the CDF of the invested times. Then, the mapping procedure proceeds as follows:

1. Compute the empirical CDF of the confidence values from the model 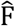 using ECDF from STATSMODELS. Trials are generated from the model such that the number of trials from each trial type follows the relative rates in data which are not uniform. The minimum number of trials generated is 10^4^.
2. Compute the empirical CDF of the wait times from data 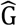 using ECDF from STATSMODELS. This is inclusive over trial types.
3. For each confidence value c, evaluate 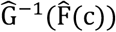. The inverse 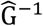 is computed via linear interpolation (using NUMPY’s interp function) inverting the x and y coordinates.

## Acknowledgments

We thank the Frank and Kepecs labs, particularly D. Liu, G. Rothschild, and T. Davidson for advice on early versions of this task; T. Ott for modeling advice; and P. Masset for the idea of temporal betting. We are grateful to J. Berke, M. Brainard, A. Nelson, and V. Sohal. We are thankful for the catalytic advice to focus on behavior and build a model, the GEMM being a product of those conversations. This work was funded by NIMH F30MH115582 (H.R.J.), F30MH109292 (J.E.C.), R01MH097061 (A.K.), NINDS, U01 NS107667 (L.M.F.), U01 NS094288 (L.M.F., A.K.); NIGMS MSTP grant T32GM007618 (J.E.C., H.R.J.); Howard Hughes Medical Institute (L.M.F.); DOE DE-AC02-05CH11231 (B.P.N). We thank NVIDIA for providing Volta GPUs for the neural network training.

## Author contributions

Conceptualization - HRJ, JEC, AK, LMF; Methodology - HRJ, JF, JEC, HL, CGB; Software - HRJ, JF, JEC, HL, BPN, CGB; Formal analysis - HRJ, HL, BPN; Investigation - HRJ, HL, CGB; Resources - LMF, AK; Data curation - HRJ, HL, JF, JEC; Writing, original draft - HRJ, HL; Writing, reviewing and editing - HRJ, HL, LMF, JF, AK, BPN; Visualization - HRJ, HL, BPN; Supervision - AK, LMF; Project administration - HRJ; Funding acquisition - LMF, AK, JEC, HRJ.

