## Supplementary Information for "Total recall: episodic memory retrieval, choice, and memory confidence in the rat"

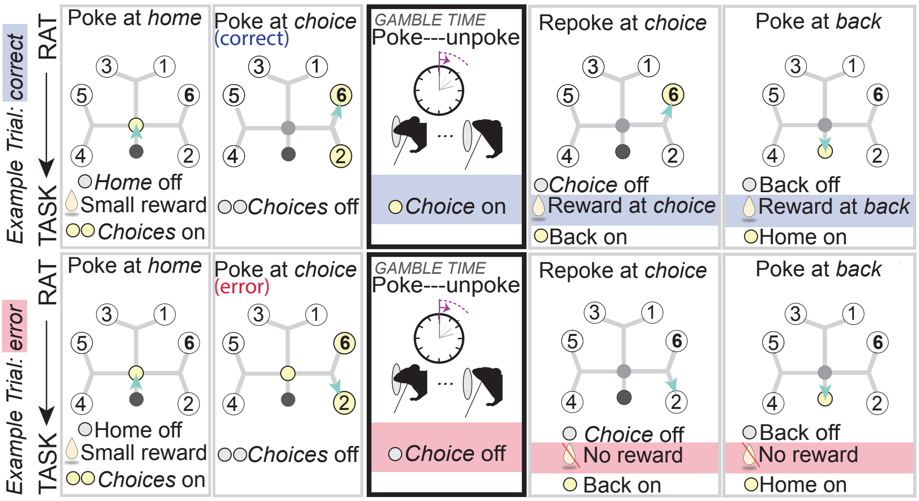


Supplementary Fig. 1. **Example trial, correct (top) and error (bottom).** Trial sequence (left to right) with rat action at top and task response at bottom of each panel. Highlighted text indicates events that occur only on correct trials (blue) or error trials (red); events that occur regardless of outcome are not highlighted. **Panel 1, top and bottom:** trials begin with rat (turquoise triangle) at home port (center, yellow), where he receives a small, fixed reward. Following a variable delay, target and distractor choice ports are cued with lights. **Panel 2:** The rat runs to and nose-pokes at his port of choice, target (port E, age 6, bold) on a correct trial (top) or distractor (port F, age 2) on an error trial (bottom). For both, the choice lights turn off. **Panel 3:** at the choice port, the rat maintains the nose-poke position for a variable, self-determined duration interpretable as the rat's confidence that the choice outcome will be correct. When he withdraws, the choice port re-lights if he was correct (top) but stays unlit if he was in error (bottom). Note that on error trials, re-poking at the choice port is not required for the task to progress, but rats did this on most trials. **Panel 4:** The rat re-pokes at the same choice port, and its light turns off and delivers reward on a correct trial (top) or no reward (bottom). The back port light turns on. **Panel 5:** The rat then runs to and pokes at the back port, which turns off, and the rat receives a reward amount equal to that received at the choice port. For an error trial (bottom), this is no reward.

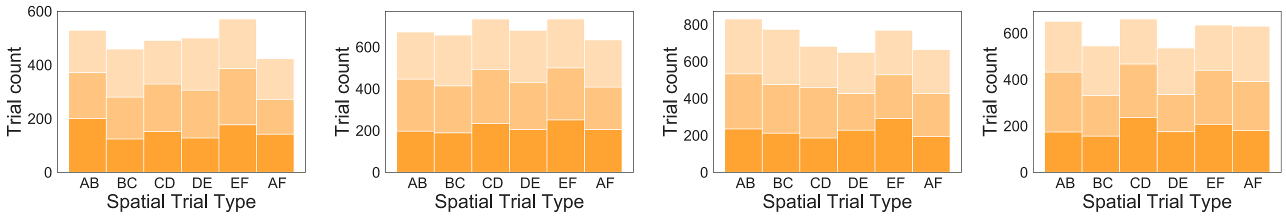


(a) (b) (c) (d)

Supplementary Fig. 2. **Even sampling of distractor ages across spatial cue pairs.** (**a-d**) From left to right, rats T, S, D, R ($n$ trials = 2978, 4111, 4369, 3660, respectively), were presented with trials approximately equally distributed over the six spatial trial types AB, BC, CD, DE, EF, and AF, and approximately equally distributed over the three distractor ages 1 (light orange), 2 (orange), and 3 (dark orange), both for a given spatial pair and overall.

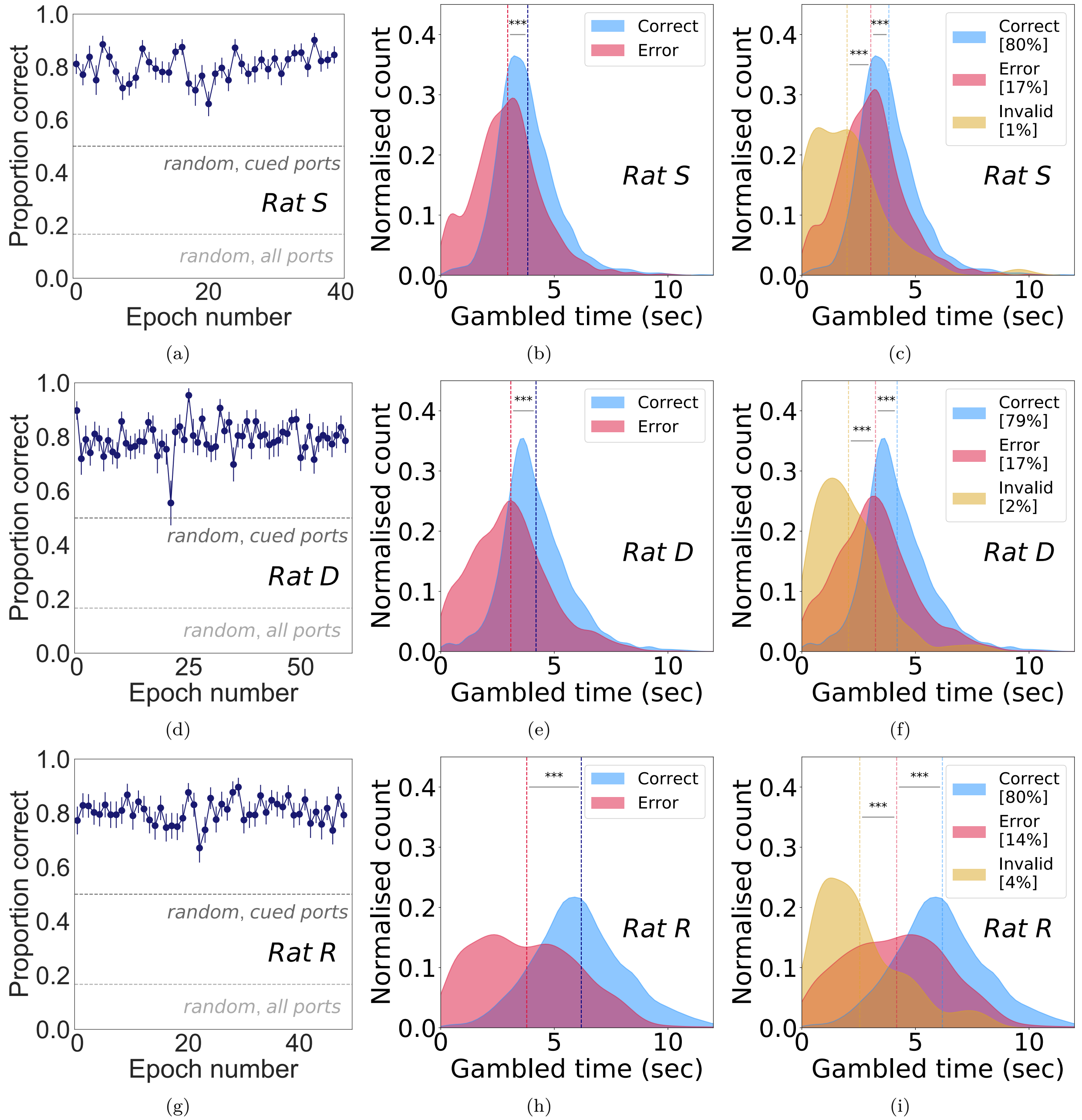


Supplementary Fig. 3. **Gambled time predicts choice accuracy for all rats.** **(a, d, g)** Choice accuracy is stable per epoch for rats S, D, R at $80.2 \pm 0.8\%, 79.4 \pm 0.7\%$, and $80.6 \pm 0.6\%$, respectively, significantly above random choice between all six ports (light gray line, 17%) or the two cued ports (dark gray line, 50%). **(b, e, h)** For all four rats, average gambled times (vertical dashed lines) for correct trials (blue) were significantly higher than error trials (red; rat S, $p = 4.9 \times 10^{-60}$; rat D, $p = 5.0 \times 10^{-81}$; rat R, $p = 6.5 \times 10^{-118}$). **(c, f, i)** For all four rats, average gambled times (dashed vertical lines) were significantly shorter for invalid choices (yellow) than for errors to the cued port (red; rat S, $p = 2.2 \times 10^{-9}$; rat D, $p = 5.6 \times 10^{-14}$; rat R, $p = 2.2 \times 10^{-17}$), which were significantly shorter than for correct choices (rat S, $p = 4.5 \times 10^{-48}$; rat D, $p = 1.6 \times 10^{-56}$; rat R, $p = 3.6 \times 10^{-71}$). All error bars represent s.e.m. and all statistical tests were one-sided rank sum.

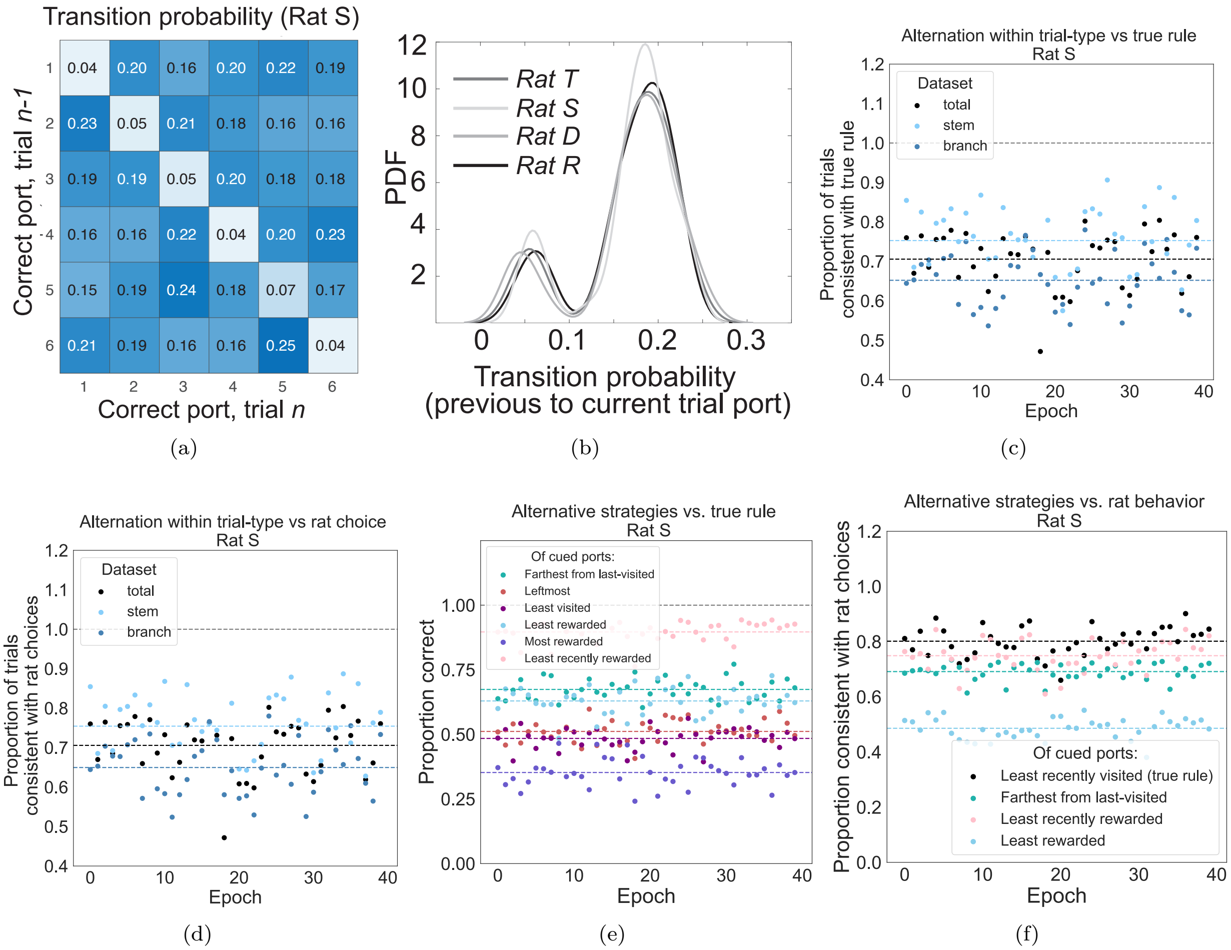


Supplementary Fig. 4. **Alternative strategies cannot explain performance accuracy.** **(a)** For all trials in all epochs from representative rat S, the correct port on trial $n-1$ (y-axis) does not strongly predict the correct port on trial $n$ (x-axis), as illustrated by the nearly equivalent transition probabilities within rows. **(b)** PDF of transition probabilities as in **a** for each of four rats shows a smooth distribution with two peaks: a low-probability peak corresponding to the same port being presented as target twice in a row, and a peak centered at 0.2, corresponding to the probability of transition from the given port to any of the other five ports. **(c)** There is no epoch (x-axis) for which an alternation rule matches the true rule on every trial (black points), stem trials only (light blue) or branch trials only (dark blue). Dashed lines = averages, all epochs. **(d)** As in **c**, for agreement with rat choices rather than true rule. There is no epoch for which an alternation strategy can explain greater than 90 percent of the rat's choices. **(e)** Alternative strategies are not consistent with the true rule. The proportion of trials that would be correct under application of each of six alternative strategies (y-axis) is shown for each epoch (x-axis) for representative rat S. The alternative strategies considered included selecting from the two cued ports the one that was: farthest from the last-visited; the leftmost; the least visited; the least cumulatively rewarded over the epoch in total; the most cumulatively rewarded over the epoch in total; or the least recently rewarded. Strategies that resulted in a correct outcome greater than 0.5 of the time were to select the least recently rewarded, the farthest from last-visited, or least rewarded overall of the two cued ports. Dashed lines = averages across epochs. **(f)** Considering for representative rat S those strategies that were in agreement with the true rule more than 0.5 of the time (blue, green, and pink points), the proportion of trials consistent with the rat's choices (y-axis) for each epoch (x-axis). The true rule (black points) explains behavior better than the alternative strategies. Dashed lines = averages across epochs.

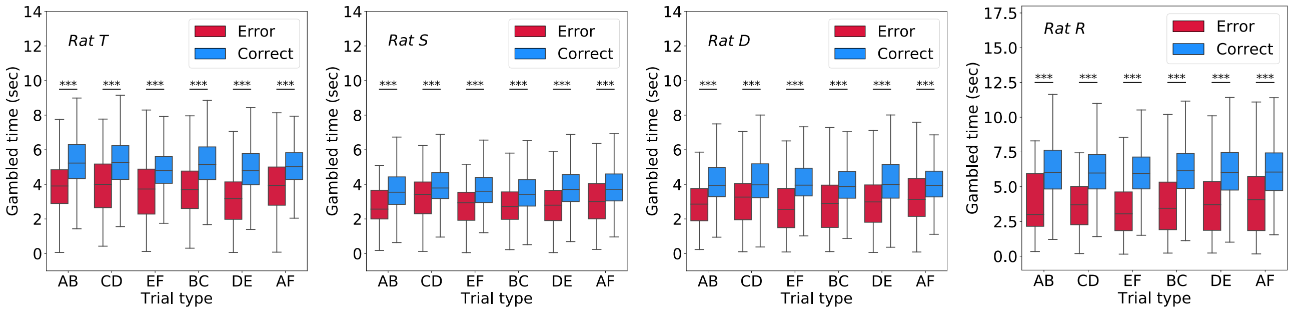


(a) (b) (c) (d)

Supplementary Fig. 5. **Gambled times are higher for correct than error trials for every port pair.** **(a)** For Rat T, pair AB, $p = 6.9 \times 10^{-9}$; pair CD, $p = 2.5 \times 10^{-9}$; pair EF, $p = 1.0 \times 10^{-7}$; pair BC, $p = 5.9 \times 10^{-17}$; pair DE, $p = 5.3 \times 10^{-23}$; pair AF, $p = 3.3 \times 10^{-9}$. **(b)** For Rat S, pair AB, $p = 9.5 \times 10^{-8}$; pair CD, $p = 1.9 \times 10^{-6}$; pair EF, $p = 1.5 \times 10^{-10}$; pair BC, $p = 3.0 \times 10^{-11}$; pair DE, $p = 5.5 \times 10^{-19}$; pair AF, $p = 8.6 \times 10^{-12}$. **(c)** For Rat D, pair AB, $p = 5.9 \times 10^{-15}$; pair CD, $p = 1.2 \times 10^{-6}$; pair EF, $p = 5.3 \times 10^{-14}$; pair BC, $p = 5.1 \times 10^{-20}$; pair DE, $p = 2.5 \times 10^{-18}$; pair AF, $p = 4.4 \times 10^{-12}$. **(d)** For Rat R, pair AB, $p = 1.1 \times 10^{-13}$; pair CD, $p = 4.5 \times 10^{-15}$; pair EF, $p = 6.6 \times 10^{-20}$; pair BC, $p = 6.5 \times 10^{-29}$; pair DE, $p = 4.8 \times 10^{-22}$; pair AF, $p = 2.0 \times 10^{-22}$. All tests are one-sided rank sum tests.

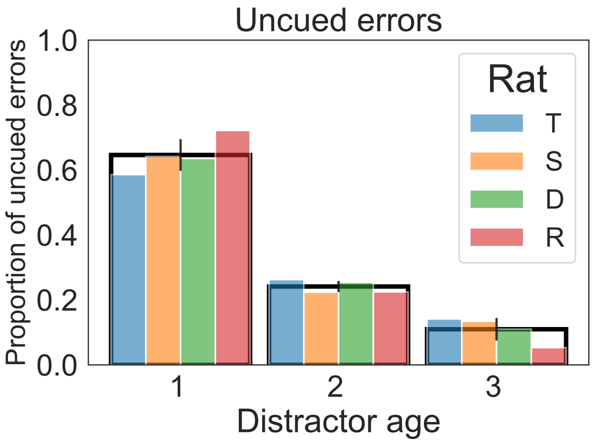


Supplementary Fig. 6. **Invalid choices were most common on trials presenting distractor age 1**. The percentage of invalid choices that occurred on distractor ages 1, 2, and 3 were: Rat T, 58.6%, 26.2%, 14.1%; Rat S, 64.2%, 22.4%, 13.4%; Rat D, 63.6%, 25.4%, 11.0%; Rat R, 72.1%, 22.5%, 5.3%.

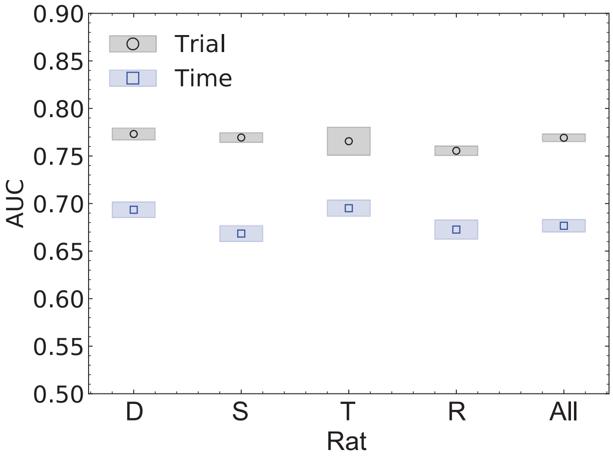


(a)

Supplementary Fig. 7. **Elapsed trials predict choice accuracy better than elapsed time.** Performance (ROC AUC) comparison of logistic regression model trained to predict choice outcome (correct/error) on target age and distractor age, last dwell time and last reward amount at each of target and distractor, the spatial relationship between target and distractor (branch/stem), and the interaction between this term and each of target age and distractor age, with target and distractor age parametrized either by clock time or number of trials. Error bars = s.e.m. across k-folds.

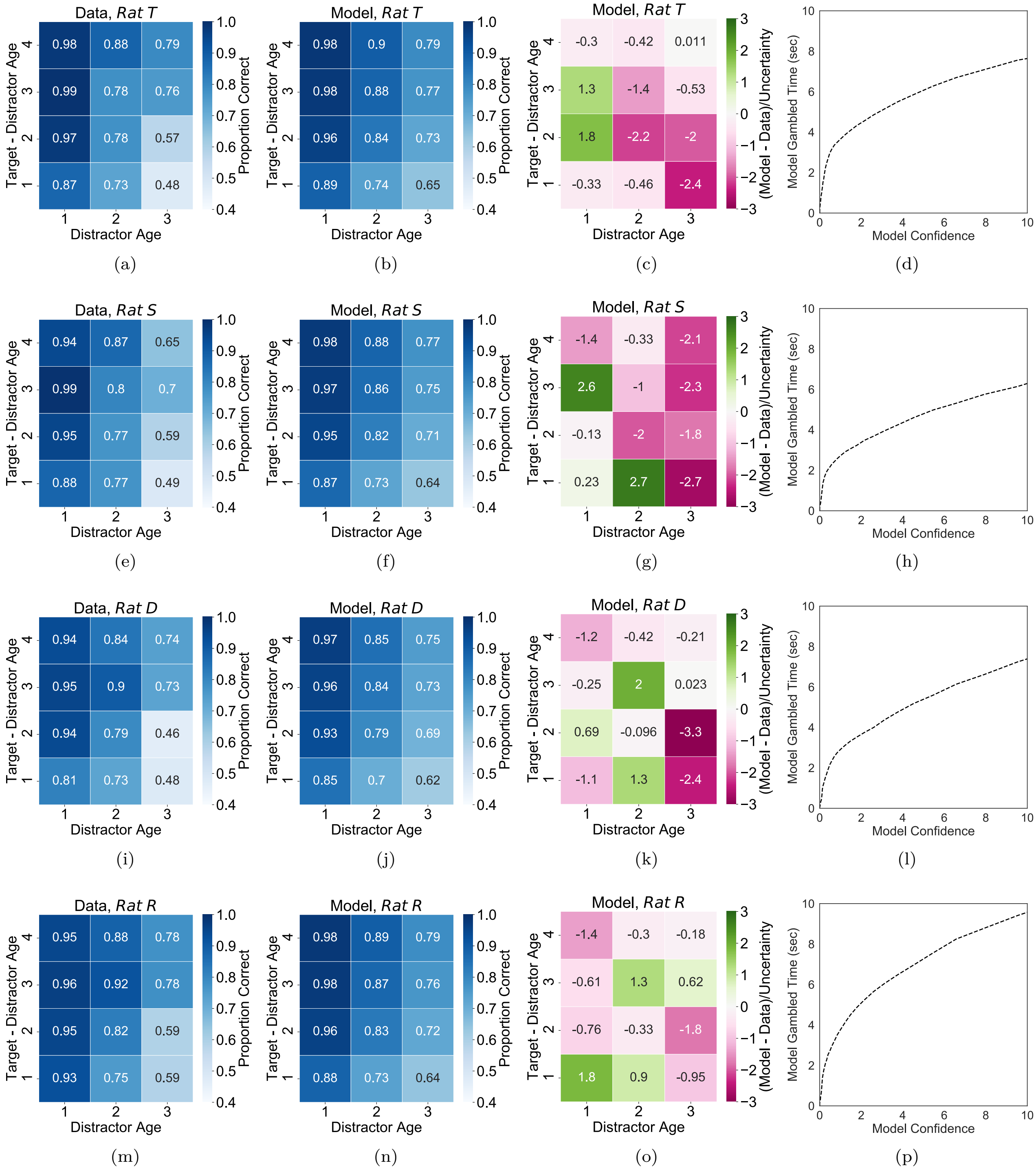


Supplementary Fig. 8. **GEMM model parameters are fit to accuracy data for each rat. (a, e, i, m)** The measured choice accuracy in bins of Distractor Age and Target – Distractor age. **(b, f, j, n)** Fitted model predictions. Fitted parameters for Rat T (**b**) were $a_0 = 1.37, a_1 = 0.33, a_2 = 0.37$, and $\sigma_0 = 0.31$. Fitted parameters for Rat S (**f**) were $a_0 = 1.29, a_1 = 0.34, a_2 = 0.35$, and $\sigma_0 = 0.33$. Fitted parameters for Rat D (**j**) were $a_0 = 1.16, a_1 = 0.46, a_2 = 0.14$, and $\sigma_0 = 0.45$. Fitted parameters for Rat R (**n**) were $a_0 = 1.41, a_1 = 0.35, a_2 = 0.30$, and $\sigma_0 = 0.34$. **(c, g, k, o)** The relative difference between the model and the data, $(\epsilon_{\text{model}} - \epsilon_{\text{data}})/\epsilon_{\sigma_{\text{data}}}$, where error rate $\epsilon = 1 - \mathit{Accuracy}$. **(d, h, l, p)** Mapping function $\hat{G}^{-1}(\hat{F}(c))$ from confidence to gambled time (see Methods).
